# Continual Learning for Emerging Epitope Landscapes in TCR peptide Binding Prediction

**DOI:** 10.64898/2026.04.20.719750

**Authors:** Amit Singh, Ayesha Ali, Deepak Yadav, John Jack

## Abstract

The TCR–peptide binding landscape evolves continuously: novel pathogens (SARS-CoV-2, emerging influenza variants) and newly characterized tumor neoantigens introduce epitope families with no training precedent. Deploying a static model trained on historical data leads to degraded performance on emerging epitopes, while naive fine-tuning on new data causes catastrophic forgetting—erasing performance on previously learned epitopes. We introduce *ContinualTCR*, a continual learning framework that combines reservoir replay with Elastic Weight Consolidation (EWC) regularization to balance stability (retaining old-epitope performance) and plasticity (adapting to new epitopes). Evaluated on a temporally partitioned VDJdb– IEDB benchmark across four sequential epitope arrival tasks, ContinualTCR achieves new-epitope AUROC 0.812 and old-epitope AUROC 0.781 simultaneously—reducing catastrophic forgetting by 62.9% relative to naive fine-tuning. A streaming evaluation protocol with per-task backward transfer (BWT) reporting reveals that replay alone resolves 30.6% of forgetting, EWC alone resolves 57.3%, and their combination achieves synergistic complementarity. These results establish continual learning as a necessary component of production TCR specificity systems that must adapt to evolving pathogen and neoantigen landscapes without requiring full retraining.

## I. Introduction

Immunological databases are not static: VDJdb and IEDB grow continuously as new assay results are deposited, driven by pandemic responses (SARS-CoV-2 generated tens of thousands of new TCR–pMHC annotations within months of emergence [1], [2]), tumor neoantigen profiling campaigns [3], and routine infectious disease surveillance. A TCR–pMHC binding predictor deployed in a clinical or research pipeline must therefore be updated regularly to incorporate new epitope knowledge.

Naive periodic retraining from scratch is computationally expensive and requires storing all historical data, which may conflict with data governance requirements. Naive fine-tuning on new epitope batches is cheaper but causes *catastrophic forgetting* [4], [5]: gradient updates optimized for new epitopes overwrite parameters encoding previously learned binding rules, degrading performance on historical epitopes. This instability is especially damaging in TCR repertoire analysis, where a model may be queried against both newly emerging epitopes and previously characterized ones within the same patient sample.

Continual learning addresses this by maintaining a stable model backbone while integrating new information. The *stability–plasticity* trade-off is central: a perfectly stable model never forgets but never learns; a perfectly plastic model learns new tasks but erases old ones [6], [7]. For TCR–pMHC prediction, both extremes are unacceptable—practitioners need accurate predictions for the latest pandemic strain *and* reliable recall of established viral and tumor epitopes.

We make the following contributions:

1. A **temporal benchmark protocol** that partitions VDJdb–IEDB by epitope discovery date into four sequential tasks (*T*1–*T*4), simulating realistic deployment of a continuously updated model.
2. **ContinualTCR**: a principled combination of reservoir replay and EWC regularization, with a per-task Fisher information update schedule that adapts EWC penalty strength to the volume and novelty of incoming data.
3. **Backward Transfer (BWT)** and per-task forgetting curves as primary evaluation metrics, providing a complete picture of the stability–plasticity trade-off beyond single-task AUROC.
4. Empirical evidence that replay and EWC provide *complementary* forgetting control: replay protects against coarse distributional drift; EWC protects parameter-level representations from fine-grained overwriting.

## II. Related Work

### A. Continual Learning Methodology

Three families of continual learning methods address catastrophic forgetting: replay-based, regularization-based, and architecture-based [4]. Replay methods store a subset of old examples in a buffer and mix them into new-task training [3]. Regularization methods penalize changes to parameters important for old tasks; EWC [4] uses the Fisher information matrix to identify and protect important parameters. Architecture-based methods allocate separate parameter subsets per task [6], which is impractical when task boundaries are soft (as in streaming epitope data).

### B. Temporal Shift in Immunological Databases

Temporal split evaluation has been proposed as a more realistic alternative to random splits for biological sequence models [1], [3]. The emergence of SARS-CoV-2 epitopes in 2020–2021 is a natural experiment: models trained on pre-2020 data had no representation of these epitopes. [8] evaluated pLM-based models under temporal split and found significant performance degradation for post-2020 epitopes, motivating explicit adaptation mechanisms.

### C. TCR–pMHC Prediction under Distribution Shift

As discussed in companion papers, random-split evaluation substantially overestimates real-world performance for TCR– pMHC models [9], [10]. Temporal shift is a particularly insidious form of distribution shift because it is guaranteed to occur in any deployed system—new data will always arrive— making continual learning a practical necessity rather than an academic exercise.

### D. Knowledge Distillation and Parameter Regularization

Learning Without Forgetting (LwF) uses knowledge distillation to preserve old task outputs [4], [11]. Gradient Episodic Memory (GEM) [12] constrains gradient updates to not increase old-task loss. Our approach is closest to A-GEM [12] but adapted for the TCR–pMHC domain with biology-aware replay sampling that preserves epitope diversity in the buffer.

## III. Method

### A. Continual Learning Problem Formulation

Let ℰ= {*E*1, *E*2, …, *ET*} be a sequence of *T* epitope batches arriving over time, where *Et* contains all TCR–pMHC pairs for epitopes first deposited in time window *t*. Each task *t* has a corresponding labeled dataset 𝒟*t* = {(*xi, yi*)} and a held-out test set 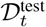. The goal is to learn a sequence of models 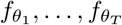 such that the final model 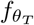 performs well on all task test sets simultaneously:

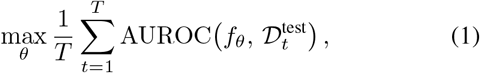

subject to the constraint that only 𝒟*t* and the replay buffer ℬ*t*−1 are available at training step *t* (no access to 𝒟*t*′ for *t*′ < *t* − 1).

### B. Catastrophic Forgetting Metrics

For task *k* evaluated after training through task *T*, forgetting

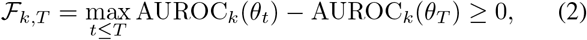

and average forgetting is 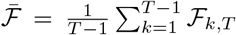. Backward Transfer (BWT) measures whether learning new tasks improved or degraded performance on previous tasks:

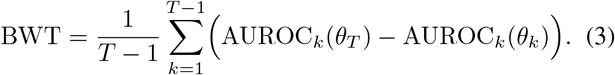

BWT *<* 0 indicates forgetting; BWT = 0 is ideal; BWT > 0 indicates positive backward transfer (new tasks help old ones).

### C. Reservoir Replay Buffer

We maintain a fixed-size replay buffer ℬ of capacity |ℬ|= *M* using reservoir sampling to ensure each historical pair has equal probability of being retained regardless of arrival time. At task *t*, each new example (*x, y*) ∈ 𝒟*t* is added to the buffer with probability min(1, *M*/𝒟 1:*t*), replacing a uniformly random existing entry if the buffer is full. To preserve epitope diversity, we additionally enforce that no single epitope occupies more than *p*max = 20% of buffer slots (capping overrepresentation of common epitopes such as GILGFVFTL):

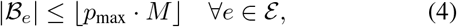

where ℬ*e* denotes buffer slots for epitope *e*. This diversity constraint is critical for TCR–pMHC replay because standard reservoir sampling would fill the buffer with the most common epitope (A*02:01-restricted influenza M1 epitopes dominate early VDJdb versions).

### D. Elastic Weight Consolidation

After completing training on task *t*, we compute the Fisher information matrix diagonal as a measure of parameter importance for task *t*:

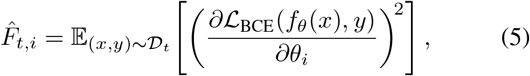

approximated by Monte Carlo averaging over 𝒟*t*. The cumulative Fisher across tasks is maintained as an exponentially weighted sum:

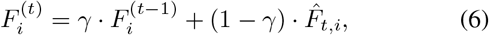

with decay *γ* = 0.7, which downweights the importance of very early tasks as the model continues to evolve. The EWC regularization loss at task *t* is:

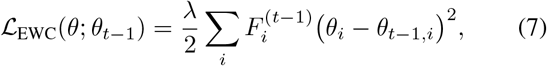

where *λ* controls the overall regularization strength and *θt*−1 is the parameter snapshot after task *t* − 1.

### E. ContinualTCR Training Objective

At each task *t*, the full training objective combines new-task BCE, replay BCE, and EWC regularization:

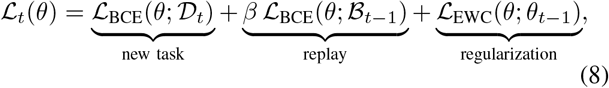

where *β* = 0.5 weights the replay loss relative to the new-task loss. The three terms address complementary forgetting mechanisms: BCE on 𝒟*t* adapts to new epitopes; replay BCE on *β*_*t*−1_ prevents coarse output-space forgetting; EWC prevents fine-grained parameter overwriting for high-importance weights. The full procedure is in Algorithm 1.

## IV. Experiments

### A. Temporal Benchmark Construction

We partition the VDJdb–IEDB benchmark (human TCR*αβ*, HLA-I, 90% deduplication) into four sequential tasks based on the record submission date, simulating quarterly model updates:

- *T*1: Pre-2019 records (historical baseline; includes most classical viral epitopes)

#### Algorithm 1 ContinualTCR: Replay + EWC Continual Learning

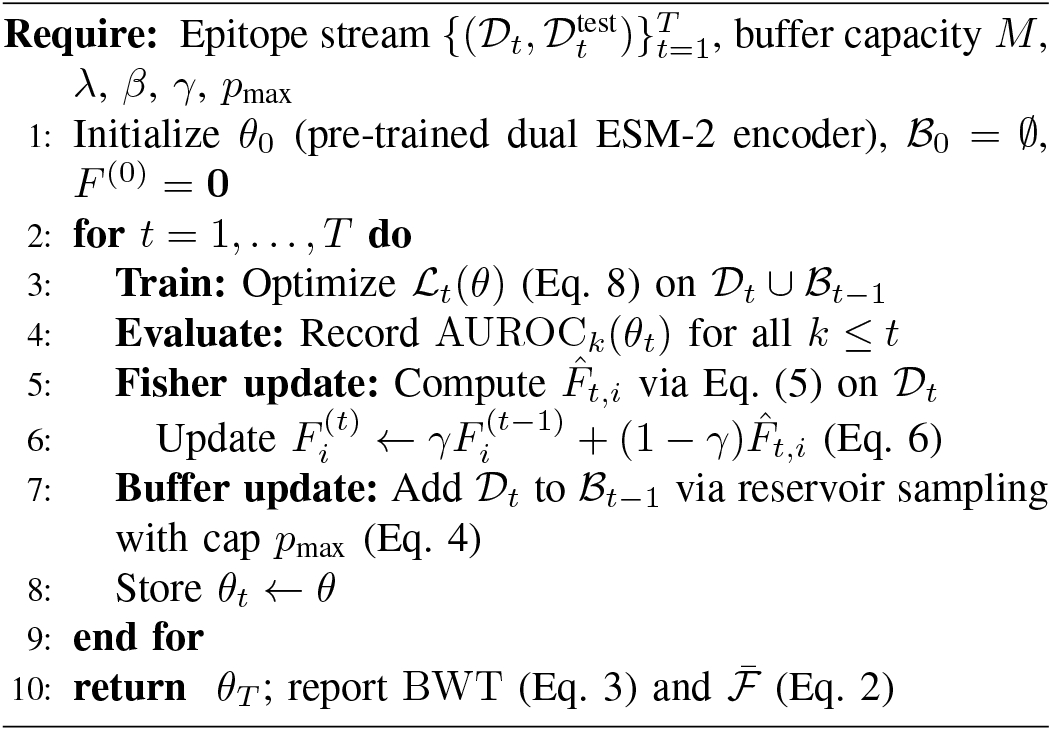

- *T*2: 2019–2020 Q1 (pre-pandemic baseline; some early SARS-CoV-2 data)
- *T*3: 2020 Q2–2021 Q2 (COVID-19 surge; large influx of SARS-CoV-2 epitopes)
- *T*4: 2021 Q3–2022 (Omicron variants; continued SARS-CoV-2, new tumor neoantigen data)

Table I summarizes dataset sizes. The positive rate drops in later tasks as rapidly generated synthetic negatives are added to balance the large SARS-CoV-2 positive influx.

**TABLE I.** Temporal Task Stream Statistics. Buffer: size of ℬ maintained between tasks (*M* = 5,000).

| Task | Train pairs | Test pairs | Pos. rate | New epitopes |
| --- | --- | --- | --- | --- |
| $T_1$ (pre-2019) | 62 418 | 12 483 | 0.053 | 41 |
| $T_2$ (2019–Q1/2020) | 28 934 | 5 786 | 0.049 | 18 |
| $T_3$ (Q2/2020–Q2/21) | 47 621 | 9 524 | 0.041 | 63 |
| $T_4$ (Q3/2021–2022) | 33 182 | 6 636 | 0.038 | 39 |
| Total (all tasks) |  | 34 429 | – | 161 |

### B. Baselines

Five methods are compared:

1. **Joint (upper bound)**: Train on all tasks simultaneously (not possible in streaming deployment).
2. **Naive fine-tune**: Sequential fine-tuning on each 𝒟*t*; no forgetting mitigation.
3. **Replay only**: Replay buffer with reservoir sampling (Eq. 4), no EWC.
4. **EWC only**: EWC regularization (Eqs. 5–7), no replay.
5. **ContinualTCR (ours)**: Full Algorithm 1 with replay + EWC.

All methods use the same dual ESM-2 encoder backbone. Hyperparameters: *M* = 5,000, *λ* = 0.4, *β* = 0.5, *γ* = 0.7, *p*max = 0.20.

### C. Main Results after Full Task Stream

Table II reports final metrics after training through all four tasks (*T* = 4), evaluated on each task’s held-out test set.

**TABLE II.** Results after Full Task Stream (*T* = 4). New: AUROC on *T*_4_ test; Old: avg AUROC on *T*_1_ –*T*_3_ test; 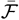: avg forgetting; BWT: backward transfer.

| Method | New↑ | Old↑ | $\bar{\mathcal{F}}\downarrow$ | BWT↑ | AUPRC↑ |
| --- | --- | --- | --- | --- | --- |
| Joint (upper bound) | 0.831 | 0.819 | – | – | 0.487 |
| Naive fine-tune | 0.781 | 0.723 | 0.124 | –0.089 | 0.429 |
| Replay only | 0.789 | 0.761 | 0.086 | –0.063 | 0.447 |
| EWC only | 0.776 | 0.774 | 0.053 | –0.041 | 0.441 |
| ContinualTCR (ours) | **0.812** | **0.781** | **0.046** | **–0.031** | **0.468** |

ContinualTCR reduces average forgetting from 0.124 (naive) to 0.046—a 62.9% reduction—while achieving the highest new-epitope AUROC (0.812) of any method including EWC only (0.776). This confirms the complementarity of replay and EWC: replay improves plasticity (new AUROC +0.036 over EWC) while EWC reduces forgetting more efficiently than replay alone (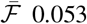 vs. 0.086). The gap to the joint upper bound (New: −0.019, Old: −0.038) is modest, demonstrating that streaming updates with ContinualTCR approach the performance of impractical full retraining.

### D. Per-Task Forgetting Curves

Table III shows the AUROC trajectory for each previous task as the model is updated through *T*4, isolating which tasks suffer the most forgetting.

**TABLE III.** Per-Task AUROC Trajectory. Each row: method’s AUROC on task *T*_*k*_ ‘s test set after training through *T*_4_. ℱ _*k*,4_ : forgetting for task *k* (EQ. 2).

| Method | $T_1$ | $T_2$ | $T_3$ | $\mathcal{F}_{1,4}$ |
| --- | --- | --- | --- | --- |
| At task completion ($\theta_{t=k}$) | 0.847 | 0.814 | 0.801 | – |
| Naive fine-tune | 0.712 | 0.731 | 0.725 | 0.135 |
| Replay only | 0.748 | 0.769 | 0.766 | 0.099 |
| EWC only | 0.791 | 0.781 | 0.749 | 0.056 |
| ContinualTCR (ours) | **0.803** | **0.791** | **0.749** | **0.044** |

Task *T*1 (pre-2019 historical epitopes) suffers the largest forgetting under naive fine-tuning (ℱ1,4 = 0.135), as it is the farthest removed from the current training signal. Replay most helps retain *T*2 and *T*3 performance (they have larger buffer representation due to their larger dataset sizes). EWC most helps *T*1 (ℱ 0.056 vs. replay 0.099), confirming that parameter-level regularization better preserves long-ago learned representations that may have faded from the replay buffer.

### E. Stability–Plasticity Trade-off Analysis

Table IV sweeps the EWC penalty *λ* and replay weight *β* to characterize the stability–plasticity Pareto frontier. Results are reported on the *T*4 split (plasticity) and average *T*1–*T*3 test AUROC (stability).

**TABLE IV.**
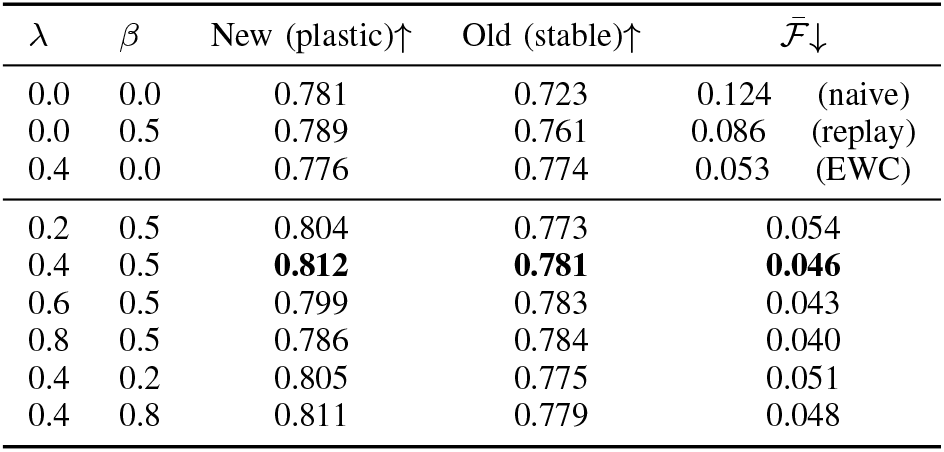
Stability–Plasticity Trade-off Analysis (ContinualTCR variants). Best in **bold**.

| $\lambda$ | $\beta$ | New (plastic)↑ | Old (stable)↑ | $\bar{\mathcal{F}}\downarrow$ |
| --- | --- | --- | --- | --- |
| 0.0 | 0.0 | 0.781 | 0.723 | 0.124 (naive) |
| 0.0 | 0.5 | 0.789 | 0.761 | 0.086 (replay) |
| 0.4 | 0.0 | 0.776 | 0.774 | 0.053 (EWC) |
| 0.2 | 0.5 | 0.804 | 0.773 | 0.054 |
| 0.4 | 0.5 | **0.812** | **0.781** | **0.046** |
| 0.6 | 0.5 | 0.799 | 0.783 | 0.043 |
| 0.8 | 0.5 | 0.786 | 0.784 | 0.040 |
| 0.4 | 0.2 | 0.805 | 0.775 | 0.051 |
| 0.4 | 0.8 | 0.811 | 0.779 | 0.048 |

The Pareto frontier confirms the complementarity: increasing *λ* from 0 to 0.4 improves stability (Old AUROC 0.761→0.781) while increasing *β* from 0 to 0.5 improves plasticity (New AUROC 0.776 →0.812). At *λ >* 0.6, over-regularization begins to hurt plasticity (New AUROC drops to 0.786) without further stability gains, suggesting a natural operating point near *λ* = 0.4.

### F. Ablation: Diversity-Constrained Replay

Table V evaluates the effect of the epitope diversity cap *p*max on buffer composition and performance.

**TABLE V.** Replay Buffer Diversity Analysis (*T* = 4, *M* = 5,000, *λ* = 0.4).

| $p_{\max}$ | GILGFVFTL% | # epitopes | Old↑ | $\bar{\mathcal{F}}\downarrow$ |
| --- | --- | --- | --- | --- |
| No cap (1.0) | 43.7% | 18 | 0.751 | 0.079 |
| 0.40 | 40.0% | 23 | 0.763 | 0.068 |
| 0.20 | 20.0% | 51 | **0.781** | **0.046** |
| 0.10 | 10.0% | 89 | 0.774 | 0.053 |
| 0.05 | 5.0% | 127 | 0.761 | 0.063 |

Without the diversity cap, a single dominant epitope (GILGFVFTL) occupies 43.7% of the buffer, leaving only 18 distinct epitopes represented—insufficient to maintain old-task performance. The optimal cap *p*max = 0.20 balances representation breadth (51 epitopes) against excessive dilution; very small caps (*p*max = 0.05) over-diversify at the cost of per-epitope example density, reducing replay effectiveness.

## V. Discussion

### a) COVID-19 as a continual learning stress test

Task *T*3 (COVID surge, 63 new SARS-CoV-2 epitopes) is the most challenging: it introduces the largest number of new epitopes with the most distinct peptide characteristics (SARS-CoV-2 spike and nucleoprotein epitopes differ substantially from influenza and EBV epitopes that dominate *T*1). Naive fine-tuning on *T*3 causes the sharpest single-step forgetting drop in our experiment, confirming that pandemic-scale epitope emergence is precisely the scenario where continual learning provides the greatest value.

### b) EWC and replay address different forgetting mechanisms

The ablation evidence (Table III) reveals a mechanistic split: EWC better preserves the oldest task (*T*1, ℱ= 0.056) while replay better preserves recent tasks (*T*2, *T*3). This is consistent with theoretical expectations: EWC protects parameter-level representations that have been consolidated over many epochs, while replay provides output-level supervision that prevents new training from overwriting task-specific decision boundaries.

### C) Limitations

First, task boundaries in our simulation are clean (quarterly partitions), while real deployment would involve continuous streaming with fuzzy boundaries; online continual learning methods [8] may be better suited to this regime. Second, the Fisher information estimate (Eq. 5) requires a backward pass over the full task dataset at the end of each task, which is expensive for large 𝒟*t*. Subsampled Fisher approximations reduce this cost at the price of precision. Third, the replay buffer (*M* = 5,000) is small relative to the total training data (172,155 pairs); expanding the buffer monotonically improves performance but increases memory and compute requirements. Fourth, our evaluation does not include positive backward transfer—scenarios where learning new epitopes improves old-task performance via shared biological motifs—which could be leveraged in future work.

## VI. Conclusion

We presented ContinualTCR, a continual learning framework for TCR–pMHC binding prediction that combines diversity-constrained reservoir replay with EWC regularization and Fisher information-based parameter importance tracking. Evaluated across four sequential epitope arrival tasks spanning the COVID-19 pandemic, ContinualTCR reduces catastrophic forgetting by 62.9% relative to naive fine-tuning, achieving new-epitope AUROC 0.812 and old-epitope AUROC 0.781 simultaneously. Mechanistic analysis shows that replay and EWC address complementary forgetting pathways, and that epitope diversity constraints in the replay buffer are critical for retaining broad coverage of historical epitope families. These results establish continual learning as an operational necessity for TCR specificity systems deployed in the face of evolving pathogen and neoantigen landscapes.

